# Linking root-associated fungal and bacterial functions to root economics

**DOI:** 10.1101/2023.12.06.570504

**Authors:** Ran Wu, Xiaoyue Zeng, M. Luke McCormack, Christopher W. Fernandez, Yin Yang, Hui Guo, Meijie Xi, Yu Liu, Xiangbin Qi, Shuang Liang, Thomas E. Juenger, Roger T. Koide, Weile Chen

**Affiliations:** College of Life Sciences, Zhejiang University, Hangzhou, China; Center for Tree Science, The Morton Arboretum, Lisle, IL, USA; College of Arts & Sciences, Syracuse University, Syracuse, NY, USA; Tianmu Mountain National Nature Reserve Administration of Zhejiang, Lin’an, China; Department of Integrative Biology, University of Texas at Austin, Austin, TX, USA; Department of Biology, Brigham Young University, Provo, UT, USA

## Abstract

Tree roots form symbioses with soil microbes to acquire nutrients, but the relationships between root nutrient acquisition strategies and microbial community composition remain poorly understood. Here, we measured root traits and root-associated fungal and bacterial guilds in 336 trees of 52 species from a subtropical forest. We found a fungal gradient from ectomycorrhizal to saprotrophic dominance, which corresponded with a shift from organic to mineral nutrient economics. This fungal gradient was aligned with the increase of root nitrogen concentration, suggesting a linkage from simple root trait to fungal-mediated carbon-nutrient cycling. We also found that the functional composition of fungal and bacterial communities was closely correlated with host root-zone pH, which often varied among coexisting trees. Root-zone pH was independent of the common root traits, underpinning a potential new gradient in the root trait space. Our findings integrate microbial functions into the root economics framework, thereby advancing the understanding of diversity of nutrient acquisition strategies across forest trees.

## Introduction

Plant species have evolved a diversity of root forms and functions to facilitate resource uptake from a wide range of environments (1, 2). This belowground root diversity plays a critical role in regulating the structure and functioning of terrestrial ecosystems (3).

Species with different root traits are suggested to vary along a fast-slow economics spectrum, ranging from inexpensive, short-lived roots with high metabolic activity to expensive, long-lived roots with lower metabolic activity (4–7). However, roots do not act alone; they associate with soil fungi and bacteria with various functions to enhance nutrient acquisition. Thus, incorporating the functions of root-associated fungal and bacterial communities into the root economics framework may better capture the holistic diversity of belowground strategies in plant nutrient acquisition.

In particular, most plants form associations with mycorrhizal fungi to increase their access to soil resources. Linking the functions of mycorrhizal fungi with the root economics framework results in a two-dimensional trait space (8). In this root economics space, one axis represents how quickly resources are invested and returned, and the other axis describes where the nutrient resources are mainly captured, either roots or root-associated mycorrhizal fungi. The mycorrhizal dependence is then strongly and positively correlated with root diameter, explaining most of the variation in the root economics of nutrient acquisition (8).

However, this collaboration framework still requires refinement. For example, the assumed positive relationship between root diameter and mycorrhizal dependence may only occur for arbuscular mycorrhizal (AM) fungi. Thicker absorptive roots usually have a larger cortical volume (9), which permits greater colonization by AM fungi (10). Cortical volume, however, may not directly influence the colonization and nutrient uptake by ectomycorrhizal (EcM) fungi because the EcM structures are often limited to the outer layers of the roots (11,12). Additionally, AM fungi primarily absorb nutrients mineralized by saprotrophic microorganisms (13,14), whereas many EcM fungi are able to bypass the mineralization pathway and directly mine nutrients from organic sources (15). Clearly, it is important to consider the differences between AM and EcM fungi with respect to the root-fungal collaboration gradient.

A second limitation of the collaboration framework is its explicit focus on the complementary role of mycorrhizal fungi in root nutrient acquisition without considering other ecologically relevant fungal and bacterial taxa (3). Roots collaborate with a variety of fungal and bacterial taxa to facilitate nutrient uptake (16). The relative abundance of fungal and bacterial guilds across plant hosts may mediate the plant’s strategy in nutrient acquisition. For example, a higher relative abundance of nitrogen-fixing bacteria may indicate a lower dependence on soil nutrients (17). In addition, a higher relative abundance of saprotrophic fungi can enhance the mineralization of rhizosphere organic nutrients (18–21), potentially promoting a mineral nutrient economy surrounding the roots (22).

Despite the critical role of microbial communities in mediating nutrient acquisition strategies, microbial functions have been largely overlooked in trait-based economics frameworks of plant roots. However, increasing evidence suggests that root-associated microbial community composition can be partly explained by root traits (23–28). On the other hand, roots interact with soil in ways that also shape microbial communities (12, 28–30). Therefore, disentangling the roles of root traits and soil properties in influencing the relative abundance of root-associated fungal and bacterial guilds may make it possible to explicitly incorporate microbial functions into the root economics framework of nutrient uptake.

Here, we surveyed the functional guilds of fungal and bacterial communities in the rhizosphere (root adherent soil) and within the absorptive roots (1st-3rd orders, 31) across 336 tree hosts of 52 species from a subtropical forest (Table S1). These trees were distributed along a nearly 1000-m elevational gradient and formed either AM or EcM symbioses. Our goal was to integrate the functions of root-associated microbial communities into the existing root economics space across a broad range of plant species.

## Results

### The root-associated fungal and bacterial communities

We identified 2,037 fungal and 2,348 bacterial ASVs in the rhizosphere soil, with similar ASV richness in the root tissue. A higher proportion of fungal sequences (76.7% in rhizosphere, 65.2% in roots) were annotated with a functional guild than bacterial sequences (33.2% in rhizosphere, 45.9% in roots) (Fig. 1, S1; Table S2). In both rhizosphere soil and root tissue, saprotrophic fungi were the most abundant fungal guild (mean 42.9% in rhizosphere, 30.2% in roots), followed by ectomycorrhizal (EcM) fungi (19.0% in rhizosphere, 15.9% in roots). Plant pathogenic fungi and AM fungi were relatively less abundant (Fig. 1, S1; Table S2). AM fungi accounted for only 0.7% of the rhizosphere ITS reads, while their relative abundance based on qPCR was 1.6% (Table S2). In the bacterial communities, taxa with the function of nitrogen fixation (5.1% in rhizosphere, 7.9% in roots) and nitrogen reduction (3.2% in rhizosphere, 3.1% in roots) were the dominant nitrogen-related guilds, while the most abundant carbon-related guild was the functional group of other aerobic chemoheterotrophs (Fig. 1, S1; Table S2).

**Fig. 1.**
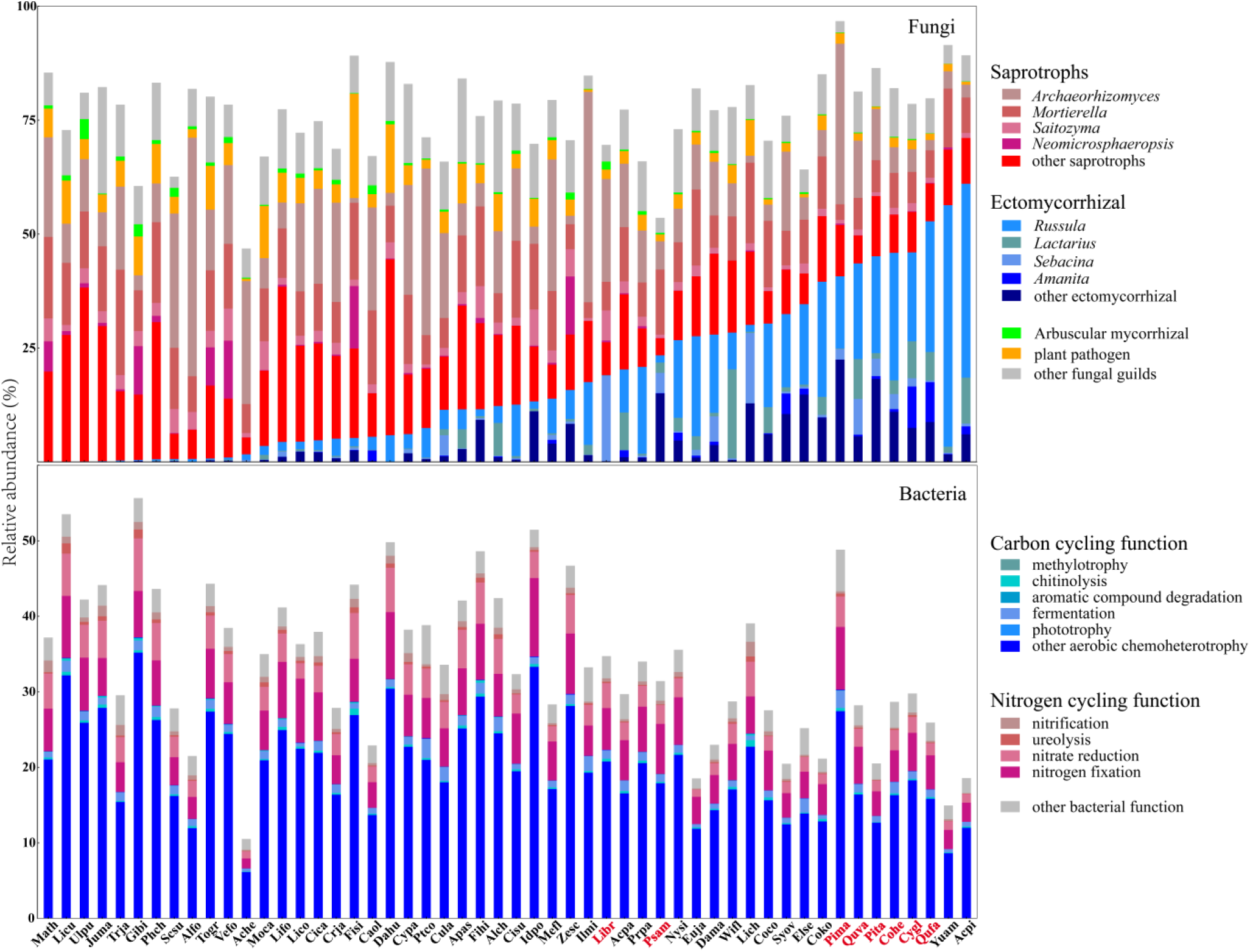
Compositions of fungal and bacterial guilds in the rhizosphere soil across 52 tree species. The relative abundance of fungal (upper panel) and bacterial (lower panel) guilds is estimated based on sequencing reads. Compositions of the five most abundant ectomycorrhizal (EcM) and saprotrophic genera are also shown. Abbreviations of EcM tree species are in red font, while arbuscular mycorrhizal (AM) tree species are in black font. Complete scientific names of trees are provided in Table S1.

### Factors influencing the compositions of fungal and bacterial guilds

Marginal tests of the dbRDA revealed that the compositions of fungal guilds in both the rhizosphere and root tissue were mainly explained by host mycorrhizal type (9.6% in rhizosphere, 13.8% in roots), elevation (2.9% in rhizosphere, 0.9% in roots), root [N] (3.0% in rhizosphere, 0.8% in roots), root-zone soil pH (5.6% in rhizosphere, 2.6% in roots) and soil gravimetric water content (1.9% in rhizosphere, 0.8% in roots) (*P* ≤ 0.05, Tables 1, S3). In contrast, the composition of bacterial guilds in both the rhizosphere and root tissue was largely unrelated to mycorrhizal type and absorptive root traits, but was strongly explained by root-zone soil pH (21.3% in rhizosphere, 13.4% in roots, Tables 1, S3).

**Table 1.** Distance-based redundancy analysis (dbRDA) to determine the predictor variables that significantly influenced the functional compositions of rhizosphere fungal and bacterial communities. Predictor variables include elevation, host mycorrhizal type, absorptive root traits, and root-zone soil properties. Results show marginal tests based on the Manhattan distance matrix, where var% indicates the relative contributions of predictor variables to fungal and bacterial guild dissimilarity. The marginal test assess each marginal term analyzed in a model with all other variables. The relative abundance of AM fungi is estimated by ITS sequencing in this table. Significant statistics (*P* < 0.05) are indicated in bold.

|  | Fungi |  |  | Bacteria |  |  |
| --- | --- | --- | --- | --- | --- | --- |
|  | var% | F | *P* | var% | F | *P* |
| Elevation | **2.9%** | **11.6** | **0.001** | **4.5%** | **19.4** | **0.001** |
| Mycorrhizal type | **9.6%** | **37.9** | **0.001** | 0.7% | 2.8 | 0.053 |
| Root diameter | 0.2% | 0.7 | 0.51 | 0.3% | 1.2 | 0.30 |
| Specific root length | 0.1% | 0.4 | 0.72 | 0.2% | 1.1 | 0.36 |
| Root [N] | **3.0%** | **11.9** | **0.001** | **1.8%** | **7.8** | **0.002** |
| Root tissue density | 0% | 0 | 1.0 | 0.4% | 1.6 | 0.18 |
| Root-zone soil pH | **5.6%** | **22.2** | **0.001** | **21.3%** | **92.7** | **0.001** |
| Soil water content | **1.9%** | **7.3** | **0.003** | 0.3% | 1.3 | 0.27 |
| Soil total carbon | 0% | 0 | 1.0 | 0.6% | 2.4 | 0.087 |
| Soil total nitrogen | 0.2% | 0.8 | 0.43 | 0.4% | 1.8 | 0.16 |
| Full model | **51.6%** | **32.3** | **0.001** | **45.2%** | **25.0** | **0.001** |

In the root economics space incorporating the guilds of fungal communities, one gradient mainly reflected the variation in root diameter, specific root length, and root tissue density. Another gradient represented all fungal guilds and root [N] (Fig.2, Tables S4-S6). In general, trees with higher root [N] tended to be associated with a greater relative abundance of saprotrophic fungi, plant pathogenic fungi, and AM fungi, but a lesser relative abundance of EcM fungi than trees with lower root [N]. This was true regardless of the sampling compartment (rhizosphere soil vs. root tissue), analysis level (individual vs. species level), or estimation method (sequencing-based or qPCR-based relative abundance of AM fungi) (Fig.S2, Tables S4-S6).

**Fig. 2.**
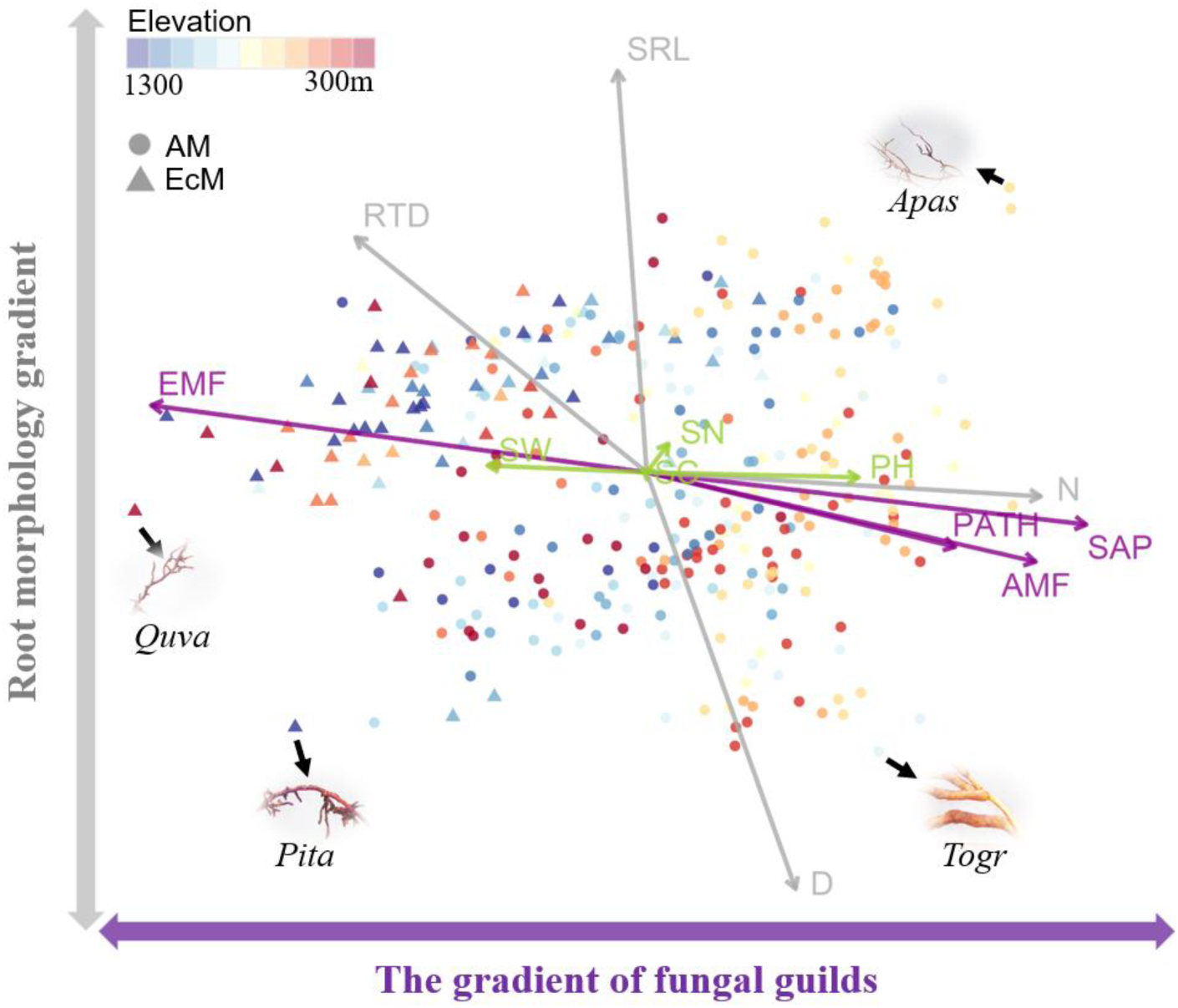
The gradient of rhizosphere fungal guilds in the root economics space. Principal component analysis followed by varimax rotation was performed on root traits (grey), root-zone soil properties (green), and relative abundances of different fungal guilds (purple). In this trait space, there is a gradient ranging from ectomycorrhizal (EcM) to saprotrophic (SAP) fungal dominance. Greater relative abundance of plant pathogenic fungi (PATH), arbuscular mycorrhizal fungi (AMF, estimated by ITS sequencing here), and higher root nitrogen concentration (N) is associated with the dominance of SAP fungi. Other root traits, including root diameter (D), specific root length (SRL), and partly root tissue density (RTD), comprise the root morphology gradient. Root-zone soil properties, including root-zone pH (PH), gravimetric water content (SW), total carbon (SC), and nitrogen (SN) concentration, mainly occupy the gradient decoupled from most root traits and fungal guilds. Four representative tree species are highlighted with pictures of absorptive roots presented at the same scale: *Quercus variabilis* (Quva); *Pinus taiwanensis* (Pita); *Torreya grandis* (Togr); *Aphananthe aspera* (Apas).

In the root economics space incorporating the guilds of bacterial communities, the gradient of bacterial guilds was strongly associated with root-zone soil pH (Figs.3, S3). Along the root-zone soil pH gradient ranging from approximately 3.5 to 7.5 across the host individuals (or from 4.0 to 6.4 at the species level), the relative abundance of rhizosphere bacterial taxa possessing the fermentation function decreased, while the relative abundance of other bacterial guilds increased (Fig.4). The remaining gradients mainly represented other root-zone soil properties, as well as host root morphology and tissue density (Tables S7-S8).

**Fig. 3.**
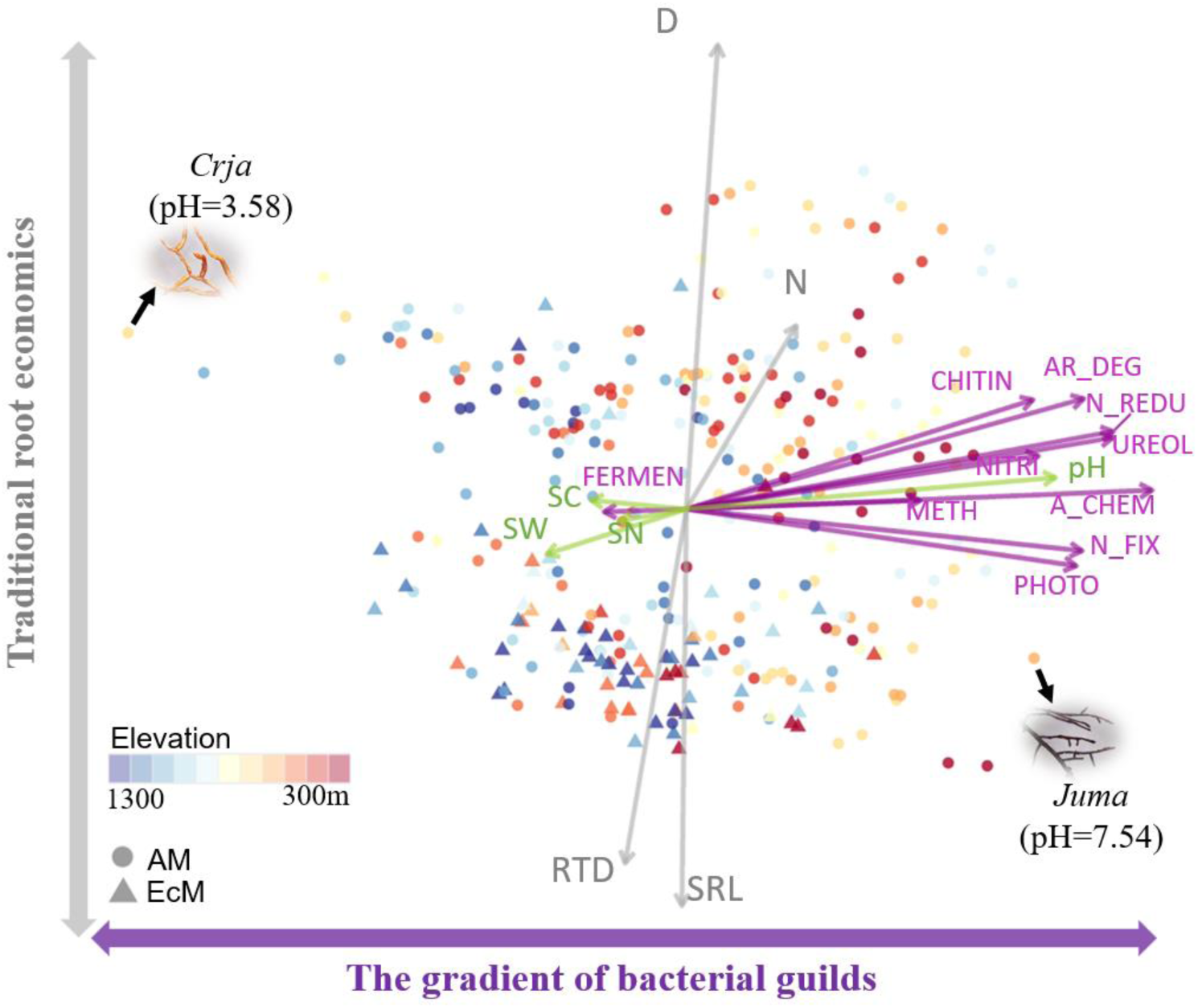
The gradient of rhizosphere bacterial guilds in the root economics space. Principal component analysis followed by varimax rotation was performed on root traits (gray), root-zone soil properties (green), and relative abundances of different bacterial guilds (purple). The variation of most bacterial guilds, including aromatic compound degradation (AR_DEG), chitinolysis (CHITIN), ureolysis (UREOL), nitrogen fixation (N_FIX), nitrification (NITRI), nitrate reduction (N_REDU), methylotrophy (METH), phototrophy (PHOTO), and other aerobic chemoheterotrophy (A_CHEM), is strongly related to root-zone pH (pH), but not gravimetric water content (SW), total carbon (SC), and nitrogen (SN) concentration. The pH gradient is also largely decoupled from the traditional root economics traits, including root diameter (D), specific root length (SRL), root nitrogen concentration (N), and root tissue density (RTD). Two representative tree species are highlighted with pictures of absorptive roots presented at the same scale: *Cryptomeria japonica* var. sinensis (Crja) and *Juglans mandshurica* (Juma).

**Fig. 4.**
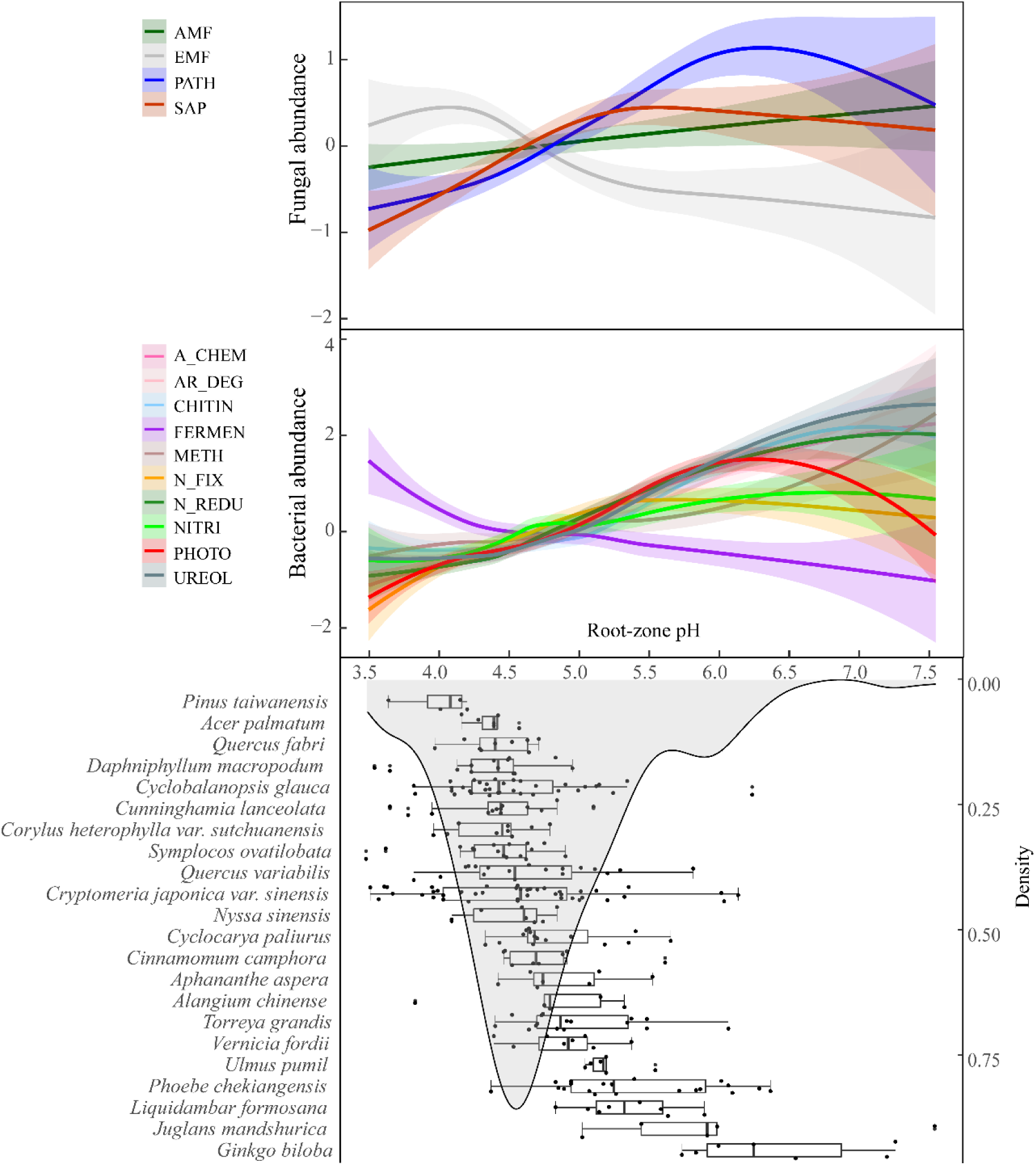
Shifts in rhizosphere fungal and bacterial functional compositions along the root-zone pH gradient. The relative abundance of different fungal (top panel) and bacterial (middle panel) guilds in the rhizosphere is standardized, plotted along the pH gradient, and fitted with a smoothed curve. The relative abundance of arbuscular mycorrhizal fungi (AMF) is estimated by ITS sequencing in this figure. In the bottom panel, root-zone pH values of the common tree species in our dataset (sample size ≥ 4) are shown (box plots), with the shaded area indicating the density curve of pH across all 336 tree hosts. All three panels share the same pH scale.

## Discussion

### The gradient of fungal guilds

When we incorporated fungal functions into the root economics space, we identified a gradient spanning from EcM to saprotrophic fungal dominance in both rhizosphere soil and root tissue of 336 subtropical trees (Fig. 2). This gradient of fungal guilds was also observed in the species-level PCAs (Fig. S2). Many EcM fungal taxa can obtain nutrients directly from organic sources (15), while saprotrophic fungi are responsible for nutrient mineralization from organic matter. Further, higher saprotrophic dominance was associated with higher proportions of AM fungi (Figs.2, S2), which enhance mineral nutrient uptake. Thus, our data clearly showed a gradient from hosts associated with fungi capable of mining organic nutrients to hosts associated with fungi that more frequently use mineralized nutrients. This gradient of root-associated fungal guilds may couple with the gradual change from organic to mineral nutrient economics across forest trees (22).

Consistent with observations in other subtropical trees, host mycorrhizal type (AM vs. EcM) was a strong factor in shaping the composition of fungal communities (Table 1; 32-33). On average, EcM trees hosted higher proportions of EcM fungi (45.1% in rhizosphere, 45.6% in roots) than AM trees (10.9% in rhizosphere, 6.1% in roots; both P <0.001 in Kruskal-Wallis rank tests). However, we detected high relative abundances of EcM fungal taxa, including *Russula* and *Lactarius* species, associated with some trees that are traditionally considered to be AM (34). Although these EcM fungi may not form typical EcM structures such as mantles or a Hartig net when associating with AM tree hosts (35), they were found in both rhizosphere soil and root tissue (Figs. 1, S1). In addition, associations between EcM fungi and conventional AM tree species have also been reported in other subtropical forests (36–37). Thus, mycorrhizal type was not the only predictor of mycorrhizal and non-mycorrhizal fungal guilds.

Indeed, root [N] explained significant variation of fungal functional composition. Higher root [N] often indicates rapid root respiration and exudation in woody species, denoting a fast root metabolic economy, whereas woody species with low root [N] typically have lower metabolic rates (6–8, 38). Thus, the trait-related root economics and fungal-driven nutrient economics may be integrated into a unified framework (Fig. 2). Moreover, our data suggest that in this integrated economics framework, the gradient of fungal guilds was largely decoupled from the axis of root morphology, suggesting an alternative gradient of fungal collaboration different from the traditional root morphology-driven framework predicting AM fungal colonization intensity (8).

This integrated economics framework largely explained the tree distribution patterns along the elevational gradient in our study site (Table 1). EcM fungi were more frequently dominant at higher elevations, where EcM trees, as well as AM trees with lower [N], were common. In contrast, saprotrophic fungi were more frequently dominant at lower elevations, where AM trees with higher [N] were more frequently distributed (Fig 2). This distribution pattern was consistent with an increase in litter decomposition rate from high to low elevations (39, 40).

The integrated economics framework was also associated with variation in the relative abundance of plant pathogenic fungi (Figs.2, S2, Tables 1, S3-S6). Colonization by EcM fungi reduces the possibility of infection by increasing pathogen resistance (41), which was supported by the negative relationships between EcM and plant pathogenic fungal abundances (Fig. 2). This negative relationship has been considered an important driver of conspecific seedling establishment within a local community dominated by EcM hosts (42, 43). Our results suggest that, in addition to mycorrhizal type, common root traits (e.g. root [N]) can also influence the abundance of EcM and plant pathogenic fungi, which may therefore explain the composition of tree species in forest communities.

### The role of root-zone pH in the root economics space

The functional composition of bacterial communities in both rhizosphere soil and root tissue was often poorly explained by common root traits, but was strongly correlated to root-zone pH (Figs.3-4, S3-S4, Tables 1, S7-S8). Along the pH gradient, low soil pH often decreases bacterial functional diversity (44) and may favor only certain bacterial guilds (e.g. fermentative bacteria in our system, Fig. S5). Also, low soil pH has been shown to inhibit chemoautotrophic growth of ammonia oxidizers, suppressing nitrification and downstream nitrogen loss pathways (45). However, the full ecological linkage between root-zone pH and bacterial functions requires further investigation.

Root-zone pH also influences the relative abundance of fungal guilds (Figs. 4, S4, Table 1). EcM fungi are often associated with more acidic soils, probably because relatively lower pH is favored by EcM fungi or because EcM fungi actively lower the pH by producing organic acids to dissolve mineral forms of phosphorus and act as chelators that are important in the oxidation of soil organic matter (46). As rhizosphere pH increases, AM fungi, saprotrophic fungi, and plant pathogenic fungi tended to become more abundant relative to EcM fungi (Fig. S4). Thus, the composition of fungal guilds was partly associated with root [N] and partly associated with root-zone soil pH (Table 1).

Importantly, root-zone soil pH was influenced by not only elevation, as previously reported (47), but also tree host identity across the study landscape, or within a plot where the local pH range was not narrow (Site 1-S6, Table S9). Therefore, in some areas, root-zone soil pH may be considered an intrinsic, measurable, tree species-specific trait. We suggest two potential causes of interspecific variation in root-zone pH. First, tree species may prefer different forms of mineral nitrogen (48), with ammonium (NH_4_^+^) uptake being associated with the release of hydrogen ions (H^+^) to the rhizosphere and lower pH, while nitrate (NO_3_^-^) uptake causes hydroxide (OH^-^) release leading to increased pH (49).

Second, tree species differ in their tissue calcium (Ca) concentration, and the Ca released by plant litter can mediate soil pH, as Ca^2+^ competes with H+ and aluminum ions (Al^3+^) for exchange sites on soil particle surfaces (50). For example, *Pinus* species often exhibit lower pH in their rhizosphere than other species, probably because of the low Ca concentration in the tissue of *Pinus* species (Fig. 4, 50).

In addition, the root-zone pH gradient was often largely decoupled from root diameter, specific root length, root [N] and root tissue density (Table S10). Although the quantity of root exudation may relate to root [N] and root tissue density (38, 51), the quality (i.e., chemical components) of the exudation may not. Only some components of the total root exudates, such as organic acids, mediate the pH (52). Thus, root-zone pH underpins a potential third gradient of the microbial-extended root economics space (Figs. 3, S3). This suggests that in addition to common root morphological and tissue chemical traits, the composition of the chemical compounds in the root exudates may also influence the rhizosphere processes through interactions with root-associated microbial communities (30).

In summary, our study has revealed new and important relationships among common root economics traits, root-zone soil properties, and the functional composition of root-associated fungal and bacterial communities under the root economics framework. First, we identified a gradient of fungal guilds spanning from EcM to saprotrophic fungal dominance. This gradient was largely decoupled from the axis of root morphology, but was strongly coupled with root [N]. Second, our findings point to the importance of the root-zone pH gradient across the tree hosts. This pH gradient was largely decoupled from the common root traits, and closely correlated with the composition of fungal and bacterial communities. These novel gradients merge the functions of absorptive roots and their associated microbial communities, resulting in a microbial-extended root economics space that can improve our ability to predict rhizosphere and ecosystem functions (e.g., organic vs. mineral nutrient economics) using simple trait-based approaches.

## Materials and Methods

### Study site and species

We collected root branches of mature trees from a subtropical forest at Tianmu Mountain, Zhejiang Province (119.4394° E, 30.3255° N). The study site has an average annual temperature of 8.8–14.8 °C and annual precipitation of 1390–1870 mm. We established eleven 50 × 50 m2 plots along an elevational gradient from 323 to 1268 m above sea level, with approximately 100-m of elevation between neighboring plots. In each plot, we sampled the roots of 6–8 of the most common tree species, with the number of tree individuals proportional to each species’ relative abundance within the plot. This resulted in a total of 336 trees sampled for this study. We queried the tree species names against The Plant List (http://www.theplantlist.org/). We assigned the mycorrhizal type of each species based on the *FungalRoot* database (53), which assumes that mycorrhizal type is usually constant within a genus. Table S1 provides the mycorrhizal type (arbuscular mycorrhizas vs. ectomycorrhizas), leaf habit (evergreen vs. deciduous), and phylogeny (gymnosperms vs. angiosperms) of each species.

### Sampling and trait measurements

We collected root and rhizosphere soil samples in early October 2020. For each tree individual, we harvested distal root branches from four random locations within 2 meters of the stem at a soil depth of 0–20 cm. We followed the target root branches to the stem to confirm the identity of the species. Each root branch included at least the first five orders (31). We collected the 1st to 3rd-order roots from each root branch as absorptive roots and pooled all absorptive roots from the root branches of the same individual tree. We stored the absorptive roots and adhering soil in Ziploc bags on ice in a cooler and transported them to the laboratory within a few hours. We then froze the samples at -80 °C for later processing.

In the laboratory, we vigorously washed absorptive root samples in 50-mL Falcon tubes using a vortex mixer for 30 seconds. We then shook the roots vigorously with sterile forceps to remove all soil from the root surfaces, and collected all root segments. We used half of the root samples to study the microbial communities associated with root tissues, which may include the rhizoplane and intraradical communities. We used the remaining half of the root samples for root trait measurements. We centrifuged the muddy solution for 2 minutes and carefully removed the supernatants with pipettes. We collected the remaining soils to study the rhizosphere microbial communities.

To measure root traits, we first pooled and scanned root segments on a desktop scanner (Epson 12000XL, Epson America, Inc., San Jose, CA, USA). We then processed the scanned images using WinRHIZO (Regent Instruments, Inc., Ottawa, ON, Canada) to determine the average root diameter and total root length. We oven-dried the scanned roots at 65 °C for 72 hours and then weighed them. We calculated specific root length as the ratio of total root length to root dry weight of the scanned roots. We calculated root tissue density by dividing the root mass by the turgid tissue volume determined by WinRHIZO. We ground the oven-dried root samples and determined the root tissue nitrogen concentration using a FLASH 2000 CHNS/O Elemental Analyzer (Thermo Fisher Scientific Inc., Waltham, MA, USA).

For each selected tree, we also sampled the surrounding soil in the field to determine the soil properties. Because the root-adherent soil was not enough for soil assays, we collected soil within approximately 5 cm of the root surface to determine the root-zone soil properties. We measured the pH of air-dried soil using an FE28 pH meter (Mettler Toledo, Shanghai, China) with a soil solution ratio of 1:2.5. We determined gravimetric water content by oven-drying at 105 °C for 72 hours. We ground the oven-dried soil and determined the carbon and nitrogen concentrations using the same elemental analyzer as for roots.

### Molecular methods

We extracted DNA from fresh root samples and root adherent soil samples using the PowerSoil DNA extraction kits (Qiagen, Germantown, MD, USA) according to the manufacturer’s instructions. After DNA extraction, we amplified the *rbcL* and *matK* gene fragments in the root DNA to verify the plant species identity (54). We amplified the ITS1 region of the fungal communities using the primer set ITS1F-ITS2 (CTTGGTCATTTAGAGGAAGTAA / GCTGCGTTCTTCATCGATGC). We amplified the V5-V7 regions of bacterial 16S rRNA using the primer set 799F-1193R (AACMGGATTAGATACCCKG / ACGTCATCCCCACCTTCC) to study the bacterial communities. We included negative controls during DNA extraction and PCR amplification.

We sent successful PCR products for high-throughput sequencing on the Illumina Novaseq platform (2 x 250 bp), with approximately 50,000 reads per sample for the 16S region and approximately 100,000 reads for the ITS region. We processed the fungal amplicon sequences using QIIME2 (55) and the bacterial amplicon sequences using USEARCH (56). We merged and quality-filtered the paired reads, and then identified the amplicon sequence variants (ASVs). We assigned the taxonomy of each sequence using the RDP classifier (57) with the UNITE databases for fungi (58) and the RDP databases for bacteria (59) at a 0.6 confidence threshold.

We used the *FungalTraits* database (34, version 1.2) to assign the functional guilds of fungal ASVs. If multiple lifestyles (i.e., guilds) were presented in the *FungalTraits* database, we used the primary lifestyle. We grouped the guilds into ectomycorrhizal (EcM) fungi, saprotrophic fungi, plant pathogenic fungi, fungi with other functions, and unspecified fungal ASVs (unknown taxonomy and unknown functions). We did not group fungal endophytes separately because their roles in plant nutrient acquisition are not fully understood. We calculated the relative abundance of different fungal guilds as the ratio of the corresponding read numbers to the total fungal read numbers.

In particular, we used two methods to estimate the relative abundance of AM fungi. First, we assigned Glomeromycotan ASVs from the ITS sequences as AM fungi and calculated the sequencing-based relative abundance across all samples (60). Second, because the ITS region may not sufficiently amplify all AM fungal taxa (61), we used qPCR methods to estimate the relative abundance of AM fungi, particularly for the rhizosphere soil samples. We estimated the qPCR-based relative abundance by quantifying the DNA copy number of AM fungi relative to the DNA copy numbers of all fungi, using primers AMG1F (ATAGGGATAGTTGGGGGCAT) and AM1 (GTTTCCCGTAAGGCGCCGAA) for AM fungi, and primers ITS5 (TCCTCCGCTTATTGATATGC) and ITS4 (GGAAGTAAAAGTCGTAACAAGG) for the whole fungal community (62). Sequencing-based and qPCR-based relative abundance of AM fungi not greater than 100% were included in the subsequent statistical analyses.

We assigned the bacterial sequencing reads to bacterial guilds using the FAPROTAX database (63). The guilds included bacterial taxa with the following functions: nitrification, nitrate reduction, nitrogen fixation, ureolysis, phototrophy, aromatic compound degradation, chitinolysis, fermentation, methylotrophy, other aerobic chemoheterotrophy (defined as aerobic chemoheterotrophy other than ligninolysis, chitinolysis, xylanolysis, cellulolysis, methanogenesis, methylotrophy, and aromatic compound degradation), bacteria with other functions (as one group), unspecified bacterial ASVs (unknown taxonomy and unknown functions). We calculated the relative abundance of different bacterial guilds as the ratio of the corresponding read numbers to the total bacterial read numbers.

### Statistical analyses

We performed fungal and bacterial-related analyses separately because the relative abundance of a fungal guild is not directly comparable to the relative abundance of a bacterial guild. We first used marginal tests of distance-based redundancy analyses (dbRDA) to determine the contributions of mycorrhizal type, plot elevation, absorptive root traits, and root-zone soil properties in shaping the composition of fungal or bacterial guilds across the rhizosphere and root tissue samples. We used the *dbrda* function in the R package vegan.

Next, we performed principal component analyses (PCAs) on the relative abundance of different fungal or bacterial guilds, as well as the corresponding root traits and root-zone soil properties across all tree individuals to construct the microbial-extended root economics space. We also performed PCAs at the species level by averaging each variable across the tree individuals of the same species. We used Horn’s parallel analysis in the R package *paran* to determine the dimensionality of these PCAs (64) and applied varimax rotation to the selected components with the principal function in the R package *psych* (65). We used varimax rotation to simplify the interpretations of the most important axes in the root economics space (66). We ln(x) transformed all data of root traits and soil properties to meet the normality requirement. To deal with zero values, we ln(x+0.001) transformed the relative abundance of fungal and bacterial guilds. We standardized the data used for principal component analyses by subtracting the mean and dividing by the standard deviation. We performed all statistics using R (version 4.1.1; R Foundation for Statistical Computing; www.r-project.org).

## Acknowledgments

### Funding

Zhejiang Provincial Funds for Distinguished Young Scientists LR21C030002 (WC)

National Natural Science Foundation of China 32101293 (WC)

National Natural Science Foundation of China 32271623 (WC)

National Natural Science Foundation of China 32001163 (SL)

National Key R&D Program of China 2022YFF1301700 (WC)

Natural Science Foundation of Zhejiang Province LQ23C030004 (RW)

Natural Science Foundation of Zhejiang Province LY23C030004 (SL)

## Author contributions

Conceptualization: WC

Methodology: WC, RW, SL

Investigation: RW, XZ, YY, HG, MX, YL, XQ

Visualization: RW, WC

Supervision: WC, SL

Writing—original draft: WC, RW

Writing—review & editing: MLM, CWF, TEJ, RTK

## Data and materials availability

All data included in this study will be available upon the acceptance of the manuscript.

## Supplementary Materials for

**This PDF file includes:**

Figs. S1 to S4

Tables S1 to S10

**Fig. S1.**
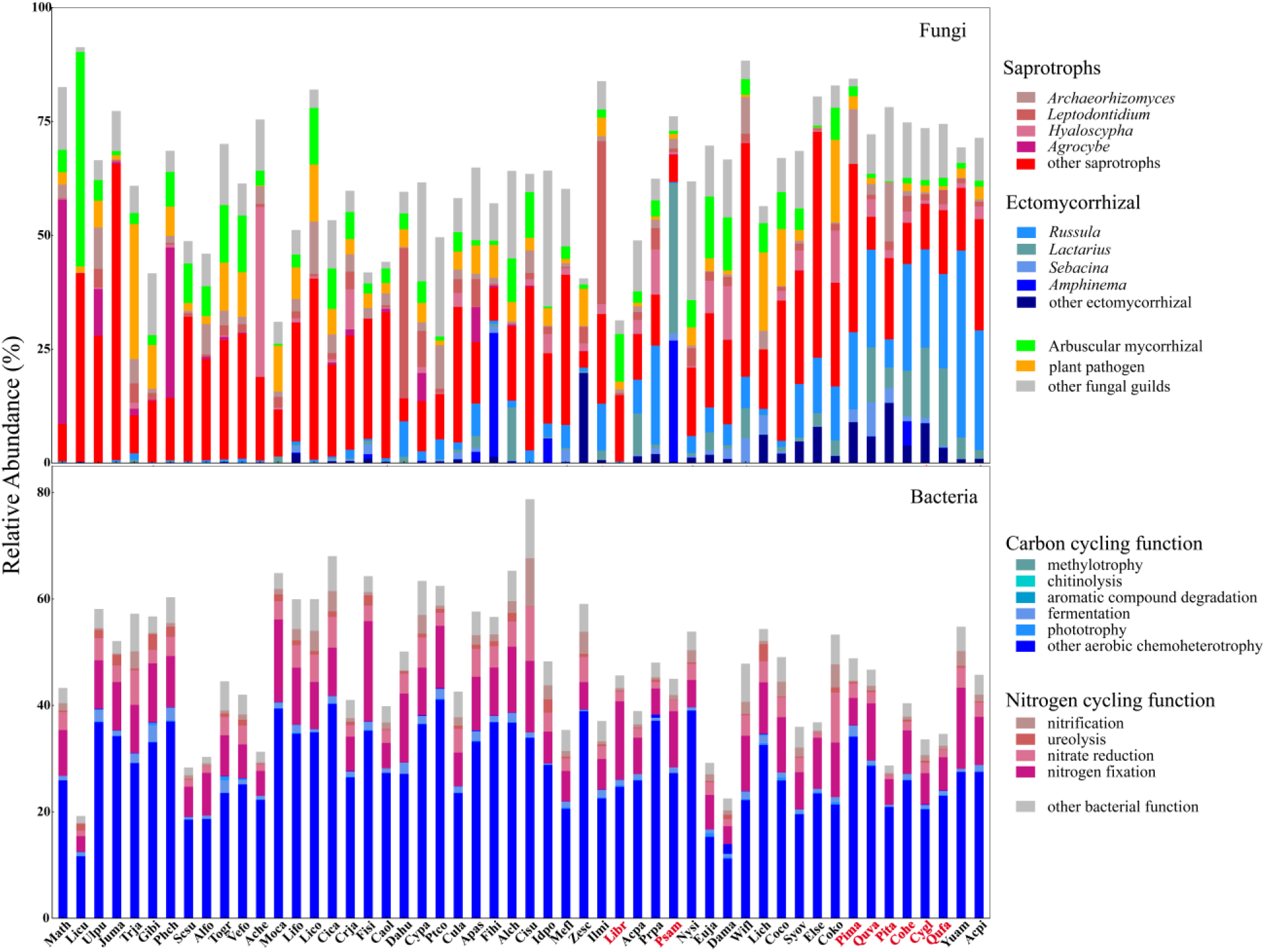
Compositions of fungal and bacterial guilds in the root tissue across 52 tree species. The relative abundance of fungal (upper panel) and bacterial (lower panel) guilds is estimated based on sequencing reads. Compositions of the five most abundant ectomycorrhizal and saprotrophic genera are also shown. Abbreviations of ectomycorrhizal tree species are in red font while arbuscular mycorrhizal tree species are in black font. Complete scientific names of trees are provided in Table S1.

**Fig. S2.**
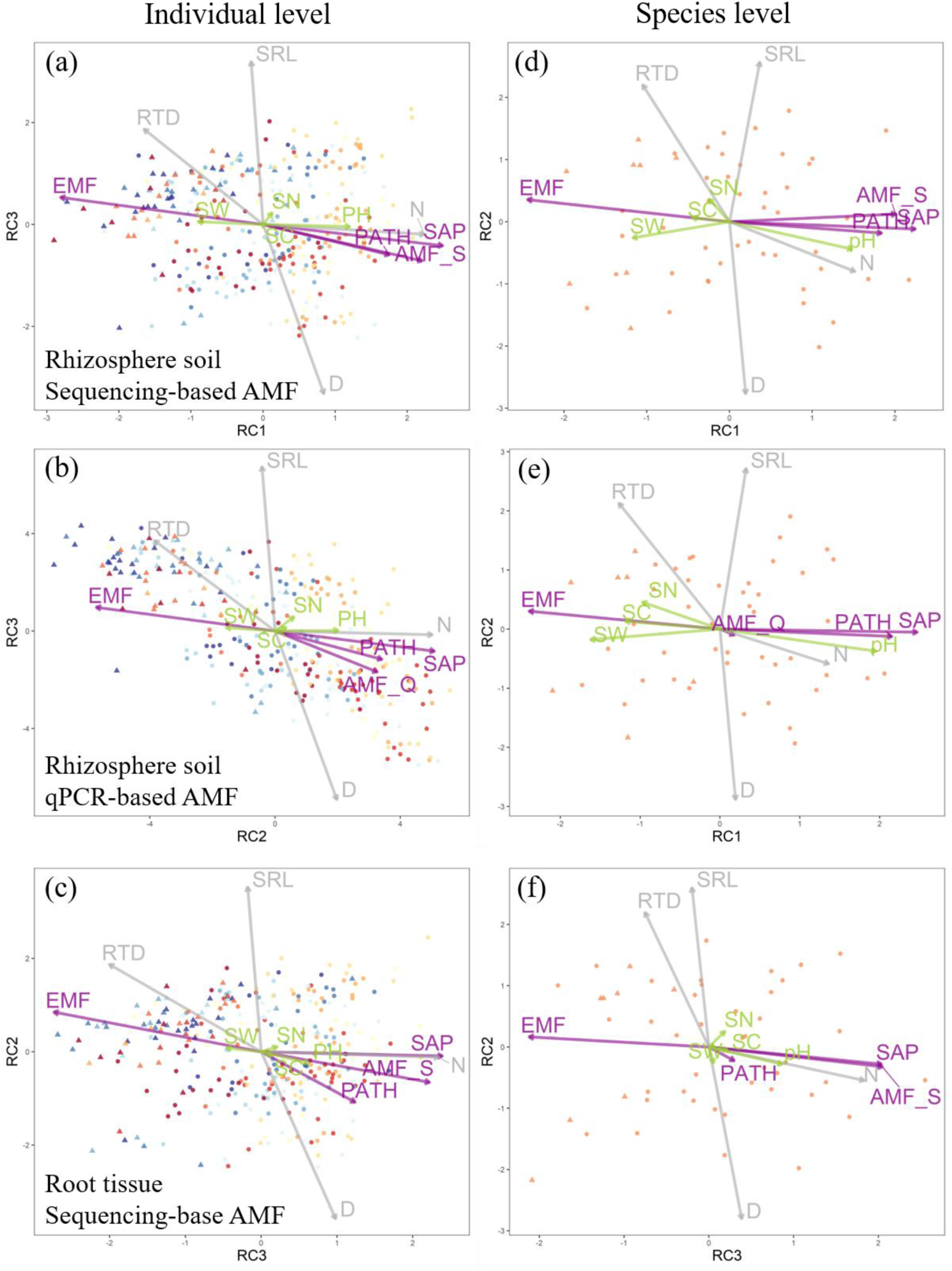
The gradient of fungal guilds (x-axis) in the root economics space. Principal component analyses followed by varimax rotation were performed based on on root traits (grey), root-zone soil properties (green) and relative abundances of different fungal guilds (purple). The data are analyzed at either the individual (a-c) or the species level (d-f). The fungal communities are sampled from either rhizosphere soil (a-b, d-e) or root tissue (c, f). Within the rhizosphere fungal communities, the relative abundance of AM fungi is estimated using either the sequencing method (a, c, d, f, AMF_S) or the qPCR method (b, e, AMF_Q). Statistics of the PCAs are shown in Tables S4-S6. See Fig. 1 for abbreviations of variables.

**Fig. S3.**
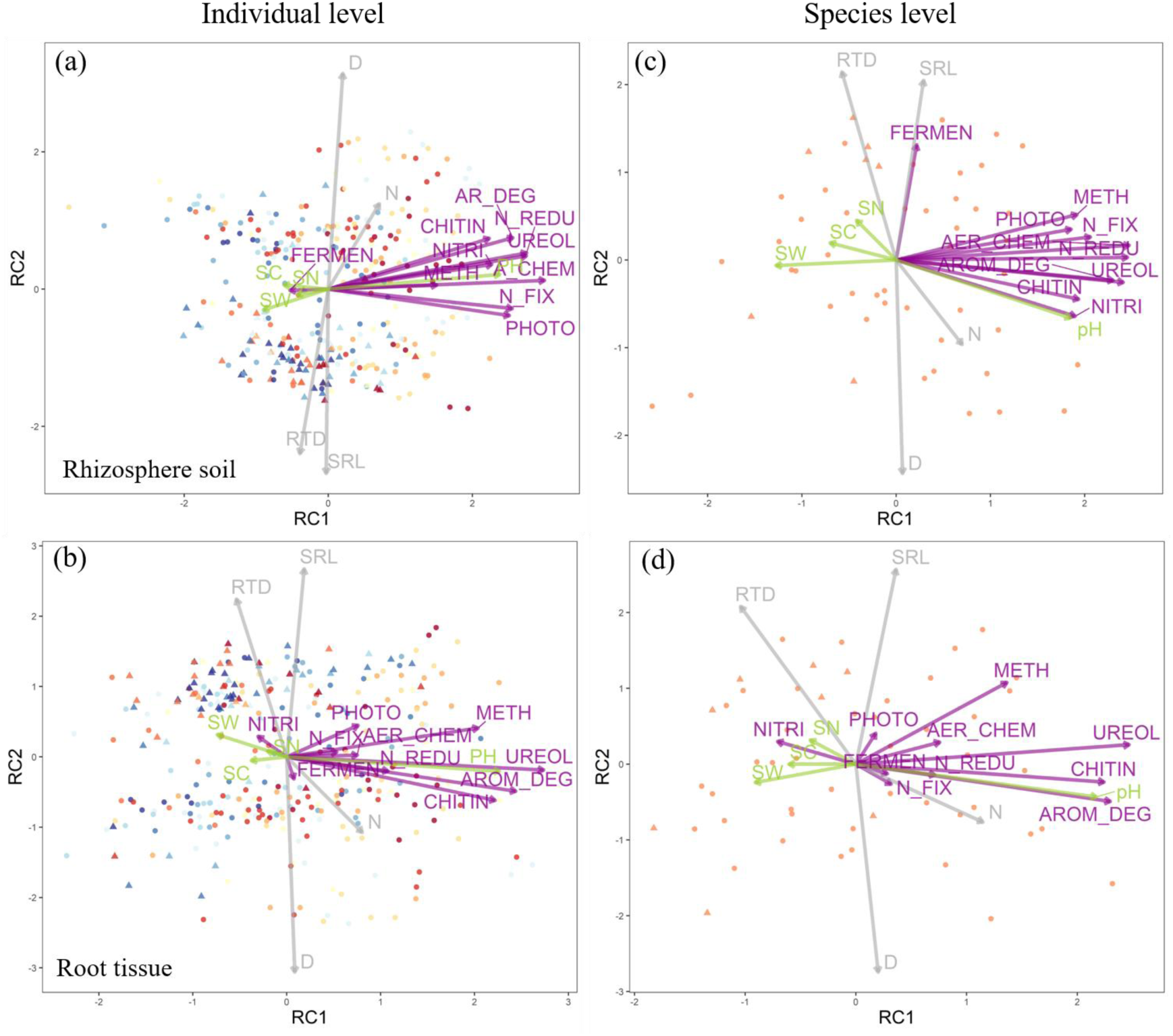
The gradient of bacterial guilds (x-axis) in the root economics space. Principal component analyses followed by varimax rotation were performed based on root traits (grey), root-zone soil properties (green) and relative abundances of different bacterial guilds (purple). The data are analyzed at either the individual (a-b) or the species level (c-d). The bacterial communities are sampled from either rhizosphere soil (a, c) or root tissue (b, d). Statistics of the PCAs are shown in Tables S7-S8. See Fig. 2 for abbreviations of variables.

**Fig. S4.**
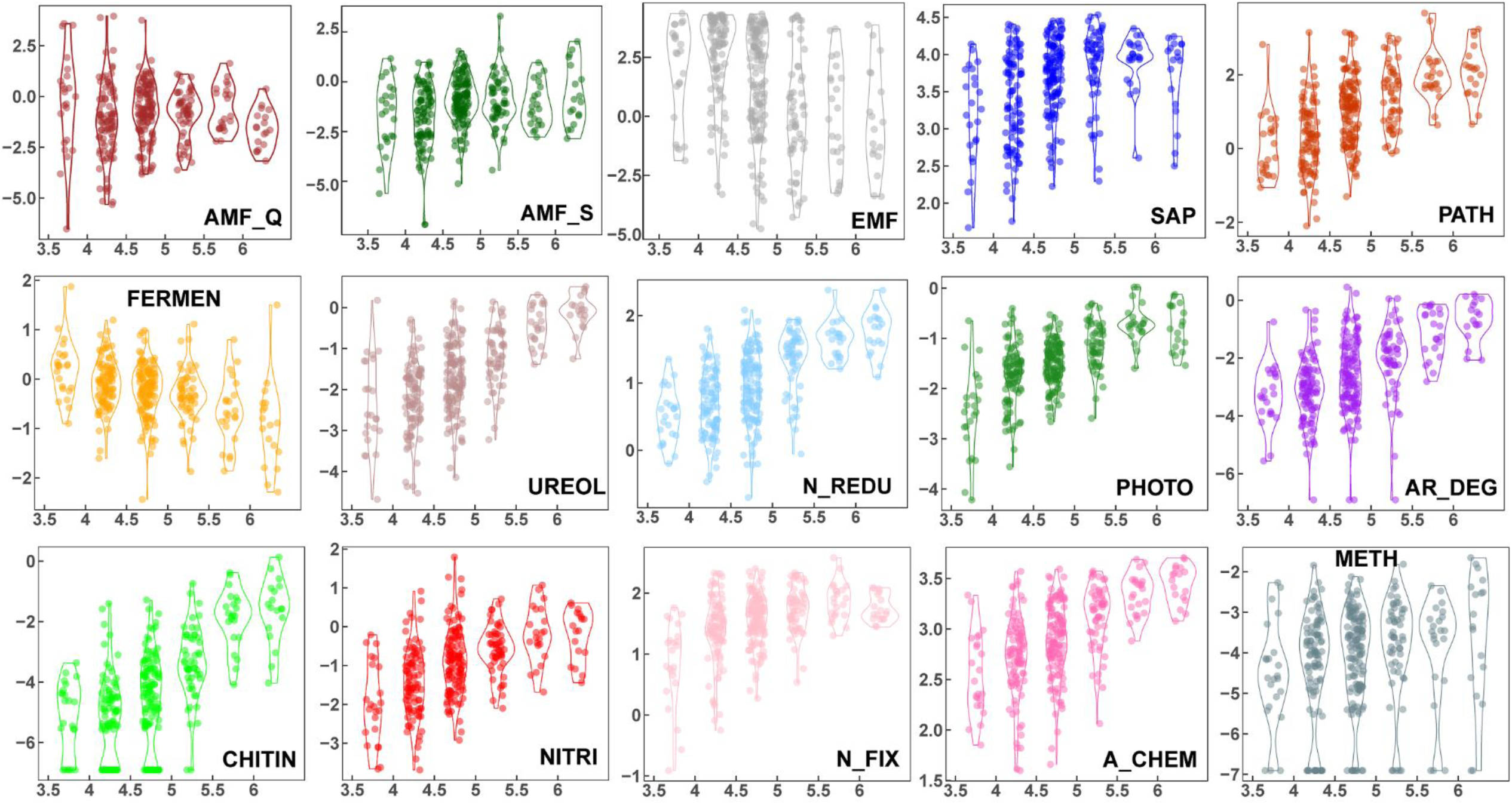
Relative abundance of different fungal and bacterial guilds along the rhizosphere pH intervals. Relative abundance of AM fungi is estimated using either the qPCR method (AMF_Q) or the sequencing method (AMF_S). Data are ln(x+0.001) transformed. See Figures 1&2 for abbreviations of fungal and bacterial guilds.

**Table S1.**
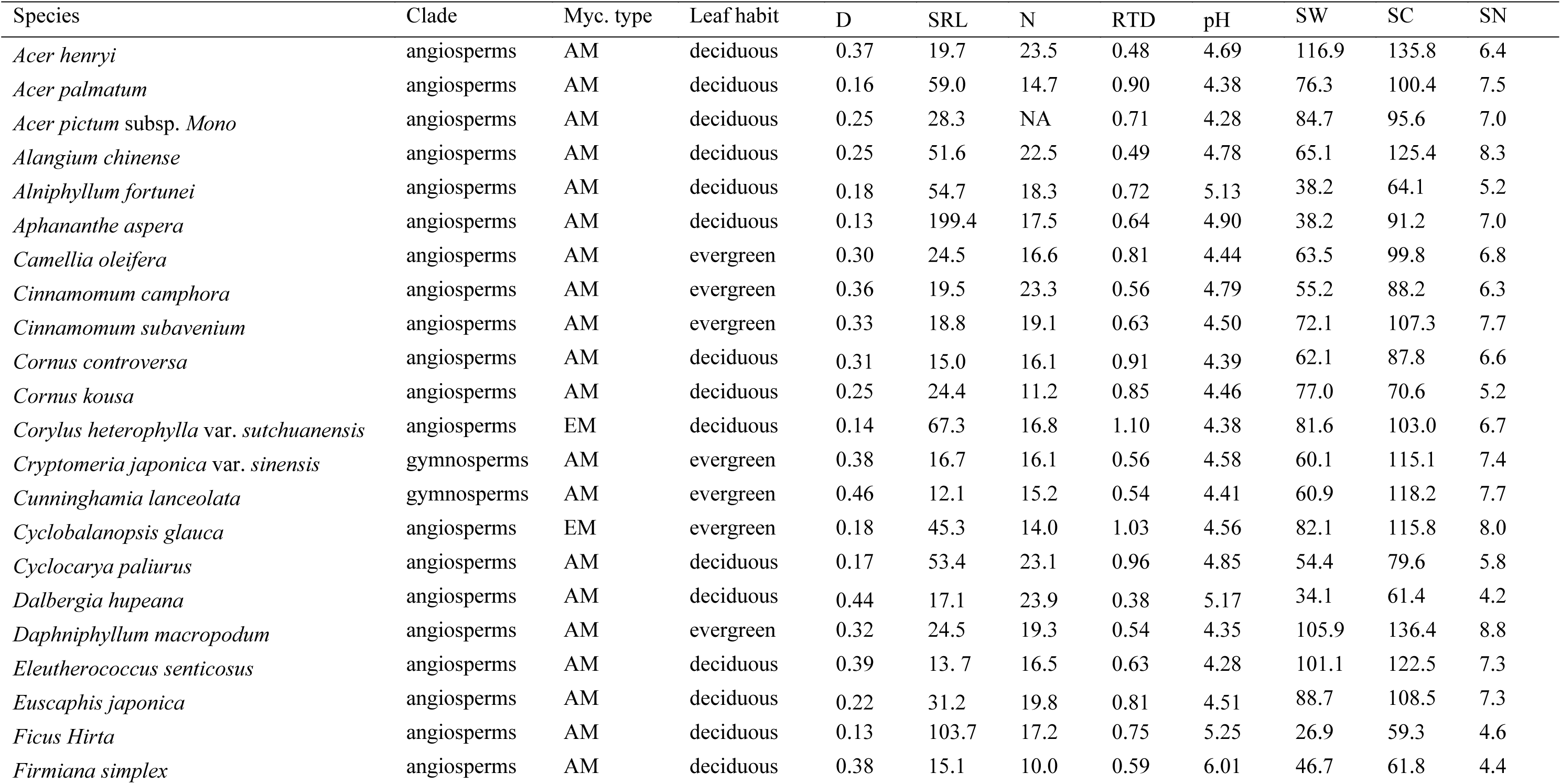
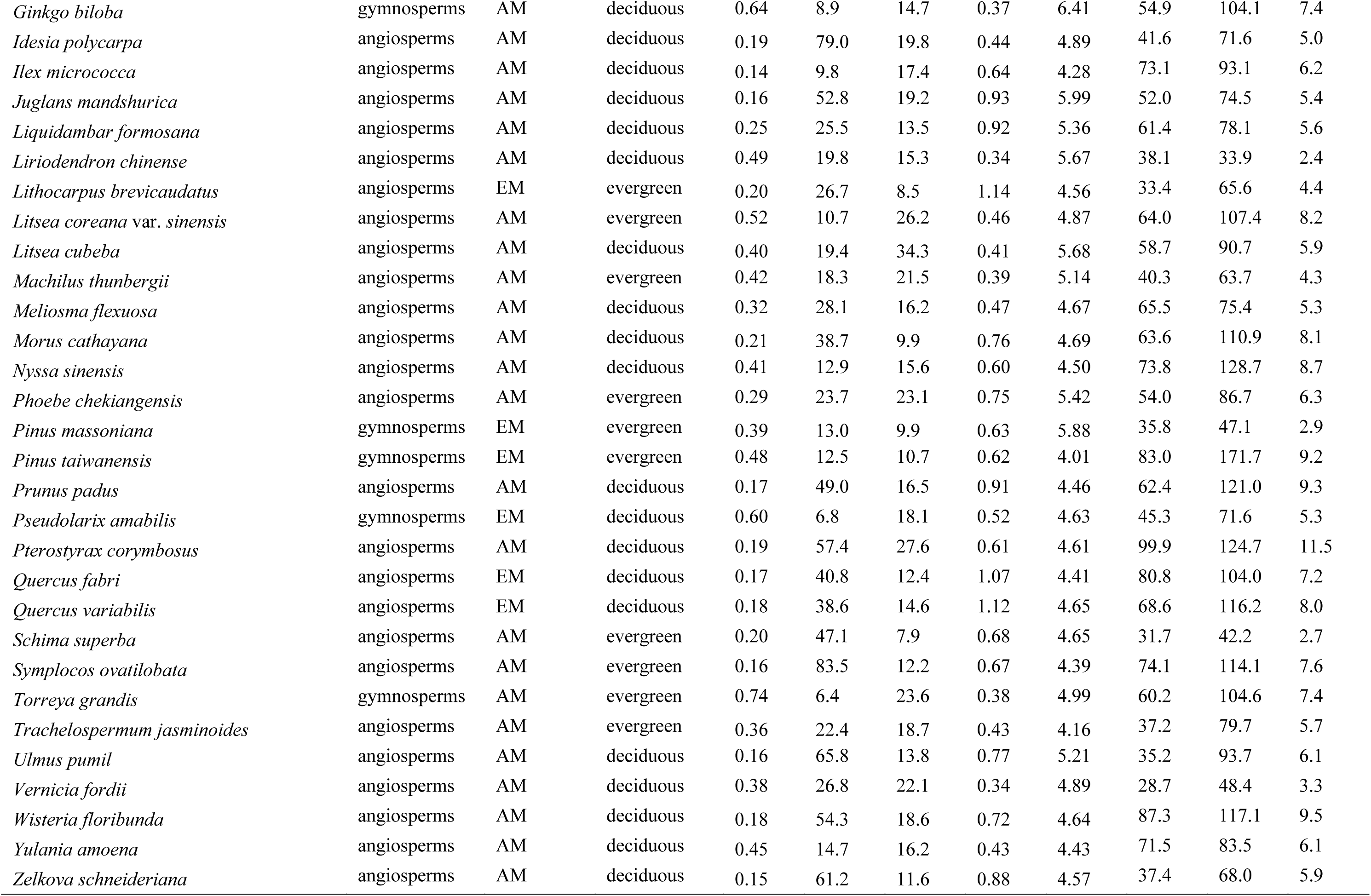
Tree species in this study. Both arbuscular mycorrhizal (AM) and ectomycorrhizal (EM) tree hosts, either evergreen or deciduous, were selected. Only the first three order roots were used for morphology and tissue chemistry measurements. Values were averaged across individuals of the same species. Root-zone soil properties were measured within ca. 5 cm of root surface. D = root diameter (mm), SRL = specific root length (m g^-1^), N = root nitrogen concentration (mg g^-1^), RTD = root tissue density (g cm^-3^), pH = root-zone soil pH, SW = gravimetric soil water content (%), SC = soil total carbon (mg g^-1^), SN = soil total nitrogen (mg g^-1^).

**Table S2.** Summary of the root-associated fungal and bacterial communities. Full names of fungal and bacterial guilds are provided in Figs 1&2. The relative abundance of AM fungi is estimated using either the qPCR method (AMF_Q) or the sequencing method (AMF_S).

|  | Rhizosphere soil |  | Root tissue |  |
| --- | --- | --- | --- | --- |
|  | Number<br>of ASV | relative<br>abundance | Number<br>of ASV | relative<br>abundance |
| Fungal guilds |  |  |  |  |
| Total | 2037 | / | 2019 | / |
| SAP | 594 | 42.87% | 589 | 30.16% |
| EMF | 257 | 18.97% | 254 | 15.88% |
| PATH | 111 | 4.37% | 109 | 4.14% |
| AMF_S | 166 | 0.66% | 166 | 5.03% |
| Others | 193 | 9.87% | 192 | 10.00% |
| AMF_Q | / | 1.58% | / | / |
| Bacterial guilds |  |  |  |  |
| Total | 2348 | / | 2352 | / |
| AR_DEG | 9 | 0.18% | 9 | 0.35% |
| CHITIN | 2 | 0.05% | 2 | 0.08% |
| FERMEN | 36 | 0.93% | 36 | 0.80% |
| UREOL | 17 | 0.30% | 17 | 0.75% |
| N_FIX | 24 | 5.13% | 24 | 7.92% |
| NITRI | 6 | 0.57% | 6 | 1.29% |
| N_REDU | 80 | 3.15% | 80 | 3.05% |
| METH | 5 | 0.03% | 5 | 0.13% |
| PHOTO | 11 | 0.27% | 11 | 0.16% |
| A_CHEM | 480 | 19.93% | 480 | 27.54% |
| others | 68 | 2.78% | 72 | 3.85% |

**Table S3.** Distance-based redundancy analysis (dbRDA) to determine the predictor variables that significantly influenced the functional compositions of fungal and bacterial communities within the roots. Predictor variables include elevation, host mycorrhizal type, absorptive root traits, and root-zone soil properties. Results show marginal tests based on the Manhattan distance matrix, where var% indicates the relative contributions of predictor variable to fungal and bacterial guild dissimilarity. The relative abundance of AM fungi is estimated by ITS sequencing in this table. Significant statistics (*P* < 0.05) are indicated in bold.

|  | Fungi |  |  | Bacteria |  |  |
| --- | --- | --- | --- | --- | --- | --- |
|  | var% | F | P | var% | F | P |
| Elevation | **0.9%** | **3.5** | **0.033** | **2.0%** | **7.3** | **0.001** |
| Mycorrhizal type | **13.8%** | **53.2** | **0.001** | 0.5% | 1.7 | 0.13 |
| Root diameter | 0.2% | 0.6 | 0.58 | 0.2% | 0.7 | 0.52 |
| Specific root length | 0.2% | 0.7 | 0.56 | 0.2% | 0.6 | 0.63 |
| Root [N] | **0.8%** | **3.0** | **0.05** | 0.6% | 2.4 | 0.067 |
| Root tissue density | 0.2% | 0.7 | 0.52 | 0.2% | 0.7 | 0.54 |
| Soil pH | **2.6%** | **9.9** | **0.001** | **13.4%** | **50.0** | **0.001** |
| Soil water content | **0.8%** | **3.0** | **0.05** | 0.2% | 0.7 | 0.56 |
| Soil total carbon | 0.7% | 2.6 | 0.07 | 0.4% | 1.4 | 0.22 |
| Soil total nitrogen | 0.2% | 0.8 | 0.47 | 0.4% | 1.4 | 0.20 |
| Full model | **40.7%** | **21.0** | **0.001** | **29.1%** | **12.5** | **0.001** |

**Table S4.** Statistics of the root economics space incorporating fungal guilds across individual trees. Shown are the proportion of variance and loadings of the relative abundance of fungal guilds, root traits, and root-zone soil properties in the most significant components of the principal component analyses followed by varimax rotation. The relative abundance of AMF is estimated by ITS sequencing. Loadings between -0.1 and 0.1 are not shown. Full names of fungal guilds, root traits, and soil properties are provided in Figure 1.

| Component | Rhizosphere soil |  |  | Root tissue |  |  |
| --- | --- | --- | --- | --- | --- | --- |
|  | RC2 | RC1 | RC3 | C1 | C2 | C3 |
| % variance | 27% | 23% | 18% | 26% | 19% | 18% |
| EMF | 0.22 | -0.82 | 0.16 | 0.23 | 0.23 | -0.73 |
| SAP | -0.31 | 0.72 | -0.12 |  |  | 0.64 |
| PATH | -0.44 | 0.51 | -0.17 | -0.48 | -0.29 | 0.33 |
| AMF | -0.21 | 0.64 | -0.21 | -0.19 | -0.18 | 0.59 |
| D |  | 0.25 | -0.96 |  | -0.95 | 0.26 |
| SRL |  |  | 0.93 |  | 0.94 |  |
| N | 0.21 | 0.65 |  | 0.12 |  | 0.62 |
| RTD |  | -0.48 | 0.54 |  | 0.50 | -0.53 |
| PH | -0.56 | 0.35 |  | -0.60 |  | 0.17 |
| SW | 0.82 | -0.26 |  | 0.85 |  | -0.12 |
| SC | 0.94 |  |  | 0.94 |  |  |
| SN | 0.94 |  |  | 0.93 |  |  |

**Table S5.** Statistics of the fungal-extended root economics space at the species level. Shown are the proportion of variance and loadings in the most significant components of the principal component analyses followed by varimax rotation. The relative abundance of fungal guilds, root traits and soil prosperities are averaged across individuals of the same species. The relative abundance of AM fungi is estimated by ITS sequencing. Loadings between -0.1 and 0.1 are not shown. Full names of fungal guilds, root traits, and soil properties are provided in Figure 1.

|  | Rhizosphere soil |  |  | Root tissue |  |  |
| --- | --- | --- | --- | --- | --- | --- |
|  | RC1 | RC3 | RC2 | RC1 | RC2 | RC3 |
| % variance | 26% | 25% | 21% | 29% | 20% | 18% |
| EMF | -0.85 | 0.11 | 0.12 | 0.34 |  | -0.73 |
| SAP | 0.78 | -0.24 |  |  |  | 0.70 |
| PATH | 0.64 | -0.32 |  | -0.61 |  | 0.10 |
| AMF | 0.70 |  |  | -0.22 | -0.11 | 0.70 |
| D |  |  | -0.96 |  | -0.96 | 0.13 |
| SRL | 0.13 |  | 0.89 |  | 0.89 |  |
| N | 0.53 | 0.46 | -0.28 | 0.22 | -0.19 | 0.63 |
| RTD | -0.37 | 0.10 | 0.77 | 0.22 | 0.76 | -0.26 |
| PH | 0.52 | -0.49 | -0.15 | -0.62 |  | 0.30 |
| SW | -0.41 | 0.80 |  | 0.90 |  |  |
| SC | -0.16 | 0.94 |  | 0.94 |  |  |
| SN |  | 0.94 | 0.13 | 0.89 |  |  |

**Table S6.** Statistics of the fungal-extended root economics space with qPCR-based relative abundance of AM fungi. Only the relative abundance of fungal guilds in the rhizosphere were included in the analyses. Shown are the proportion of variance and loadings in the most significant components of the principal component analyses followed by varimax rotation at both individual and species levels. Loadings between -0.1 and 0.1 are not shown. Full names of fungal guilds, root traits, and soil properties are provided in Figure 1.

| Fungi | Individual level |  |  | Species level |  |  |
| --- | --- | --- | --- | --- | --- | --- |
|  | RC1 | RC2 | RC3 | RC1 | RC2 | RC3 |
| % variance | 27% | 21% | 18% | 29% | 21% | 21% |
| EMF | 0.28 | -0.79 | 0.14 | -0.82 |  | 0.11 |
| SAP | -0.37 | 0.70 | -0.11 | 0.85 |  |  |
| PATH | -0.49 | 0.47 | -0.16 | 0.74 | -0.12 |  |
| AMF |  | 0.45 | -0.23 |  | 0.13 |  |
| D |  | 0.27 | -0.96 |  |  | -0.99 |
| SRL |  |  | 0.93 | 0.11 |  | 0.93 |
| N | 0.16 | 0.69 |  | 0.47 | 0.63 | -0.20 |
| RTD |  | 0.53 | 0.51 | -0.44 |  | 0.73 |
| PH | -0.60 | 0.28 |  | 0.67 | -0.31 | -0.13 |
| SW | 0.83 | -0.22 |  | -0.56 | 0.70 |  |
| SC | 0.94 |  |  | -0.41 | 0.86 |  |
| SN | 0.93 |  |  | -0.33 | 0.87 | 0.16 |

**Table S7.** Statistics of the root economics space incorporating bacterial communities across individual tree hosts. Shown are the proportion of variance and loadings of the relative abundance of bacterial guilds, root traits, and root-zone soil properties in the most significant four or five components of the principal component analyses followed by varimax rotation. Loadings between -0.1 and 0.1 are not shown. Full names of bacterial guilds, root traits, and soil properties are provided in Figure 2.

|  | Rhizosphere soil |  |  |  | Root tissue |  |  |  |  |
| --- | --- | --- | --- | --- | --- | --- | --- | --- | --- |
|  | RC1 | RC3 | RC2 | RC4 | RC1 | RC3 | RC4 | RC2 | RC5 |
| % variance | 33% | 16% | 14% | 8% | 18% | 16% | 15% | 14% | 11% |
| AR_DEG | 0.79 |  | 0.23 |  | 0.78 | -0.14 | 0.19 | -0.16 | 0.25 |
| CHITIN | 0.69 | -0.13 | 0.23 | -0.15 | 0.71 | -0.22 | 0.22 | -0.20 |  |
| FERMEN | -0.16 |  |  | 0.86 |  |  | 0.81 |  |  |
| UREOL | 0.84 | -0.16 | 0.15 |  | 0.87 |  |  |  | 0.19 |
| N_FIX | 0.78 |  |  | 0.17 | 0.17 | -0.15 | 0.79 |  | 0.34 |
| NITRI | 0.70 | -0.23 | 0.11 |  |  |  |  |  | 0.87 |
| N_REDU | 0.84 | -0.20 | 0.16 | 0.20 | 0.34 | -0.13 | 0.42 |  | 0.74 |
| METH | 0.46 |  |  | 0.57 | 0.65 |  |  | 0.13 |  |
| PHOTO | 0.77 | -0.21 | -0.12 | -0.21 | 0.24 |  | 0.75 | 0.14 | -0.19 |
| A_CHEM | 0.92 |  |  | 0.16 | 0.24 |  | 0.69 |  | 0.41 |
| D |  |  | 0.96 |  |  |  |  | -0.98 |  |
| SRL |  |  | -0.83 |  |  |  | -0.12 | 0.85 | 0.14 |
| N | 0.22 | 0.13 | 0.38 | 0.27 | 0.26 | 0.15 | -0.27 | -0.34 | 0.39 |
| RTD | -0.12 |  | -0.74 |  | -0.17 |  | 0.15 | 0.72 | -0.16 |
| PH | 0.73 | -0.28 |  | -0.41 | 0.72 | -0.37 | 0.17 |  | -0.15 |
| SW | -0.28 | 0.83 |  |  | -0.23 | 0.83 |  | 0.10 | -0.11 |
| SC | -0.19 | 0.94 |  | 0.12 | -0.12 | 0.95 |  |  |  |
| SN | -0.13 | 0.96 |  |  |  | 0.96 |  |  |  |

**Table S8.**
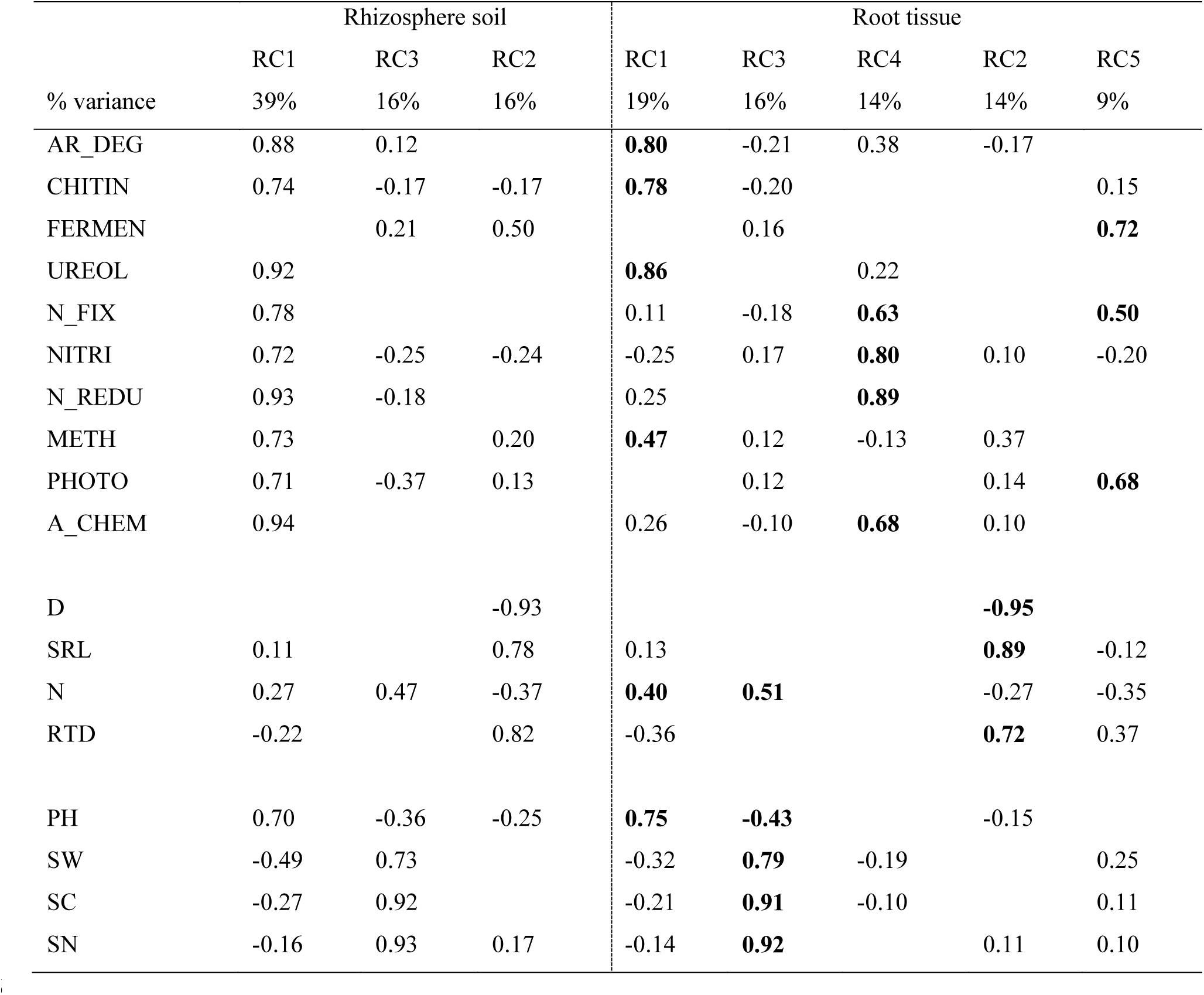
Statistics of the bacterial-extended root economics space at the species level. Shown are the proportion of variance and loadings in the most significant components of the principal component analyses followed by varimax rotation. The relative abundance of bacterial guilds, root traits and root-zone soil prosperities are averaged across individuals of the same species. Loadings between -0.1 and 0.1 are not shown. Full names of fungal guilds, root traits, and soil properties are provided in Figure 2.

|  | Rhizosphere soil |  |  | Root tissue |  |  |  |  |
| --- | --- | --- | --- | --- | --- | --- | --- | --- |
|  | RC1 | RC3 | RC2 | RC1 | RC3 | RC4 | RC2 | RC5 |
| % variance | 39% | 16% | 16% | 19% | 16% | 14% | 14% | 9% |
| AR_DEG | 0.88 | 0.12 |  | **0.80** | -0.21 | 0.38 | -0.17 |  |
| CHITIN | 0.74 | -0.17 | -0.17 | **0.78** | -0.20 |  |  | 0.15 |
| FERMEN |  | 0.21 | 0.50 |  | 0.16 |  |  | **0.72** |
| UREOL | 0.92 |  |  | **0.86** |  | 0.22 |  |  |
| N_FIX | 0.78 |  |  | 0.11 | -0.18 | **0.63** |  | **0.50** |
| NITRI | 0.72 | -0.25 | -0.24 | -0.25 | 0.17 | **0.80** | 0.10 | -0.20 |
| N_REDU | 0.93 | -0.18 |  | 0.25 |  | **0.89** |  |  |
| METH | 0.73 |  | 0.20 | **0.47** | 0.12 | -0.13 | 0.37 |  |
| PHOTO | 0.71 | -0.37 | 0.13 |  | 0.12 |  | 0.14 | **0.68** |
| A_CHEM | 0.94 |  |  | 0.26 | -0.10 | **0.68** | 0.10 |  |
| D |  |  | -0.93 |  |  |  | **-0.95** |  |
| SRL | 0.11 |  | 0.78 | 0.13 |  |  | **0.89** | -0.12 |
| N | 0.27 | 0.47 | -0.37 | **0.40** | **0.51** |  | -0.27 | -0.35 |
| RTD | -0.22 |  | 0.82 | -0.36 |  |  | **0.72** | 0.37 |
| PH | 0.70 | -0.36 | -0.25 | **0.75** | **-0.43** |  | -0.15 |  |
| SW | -0.49 | 0.73 |  | -0.32 | **0.79** | -0.19 |  | 0.25 |
| SC | -0.27 | 0.92 |  | -0.21 | **0.91** | -0.10 |  | 0.11 |
| SN | -0.16 | 0.93 | 0.17 | -0.14 | **0.92** |  | 0.11 | 0.10 |

**Table S9.** Influence of elevation and tree species on rhizosphere pH. Bold values indicate significant effects at *P*<0.05.

| Variables | F | *P* |  |  |
| --- | --- | --- | --- | --- |
| Elevation | **3.567** | **<0.001** |  |  |
| Species | **2.750** | **<0.001** |  |  |
| Elevation * Species | 1.224 | 0.15 |  |  |
| Species effect in each plot | F | *P* | pH<sub>Min</sub> | pH<sub>Max</sub> |
| S1 (323m) | **5.788** | **0.008** | 4.45 | 6.25 |
| S2 (466m) | **15.724** | **<0.001** | 3.89 | 7.26 |
| S3 (538m) | **3.284** | **0.018** | 4.16 | 7.54 |
| S4 (623m) | **13.114** | **<0.001** | 3.58 | 6.37 |
| S5 (714m) | **4.338** | **0.02** | 3.96 | 6.29 |
| S6 (809m) | **23.233** | **<0.001** | 3.79 | 7.20 |
| S7 (935m) | 1.524 | 0.25 | 3.49 | 4.89 |
| S8 (1060m) | 2.638 | 0.054 | 3.68 | 5.59 |
| S9 (1117m) | 1.199 | 0.35 | 3.83 | 5.25 |
| S10 (1190m) | 1.797 | 0.19 | 3.55 | 4.68 |
| S11 (1268m) | 0.940 | 0.45 | 3.98 | 4.64 |

**Table S10.** Principal component analysis on common root traits and root-zone soil pH. Shown are the eigenvalues, proportion of variance and loadings of the first three principal components, without loadings between -0.1 and 0.1.

|  | PC1 | PC2 | PC3 |
| --- | --- | --- | --- |
| Eigenvalue | 2.39 | 1.03 | 0.98 |
| % Variance | 47.7 | 20.6 | 19.5 |
| Diameter | 0.63 | -0.21 |  |
| Specific root length | -0.54 | 0.44 | 0.10 |
| Nitrogen concentration | 0.26 | 0.81 |  |
| Root tissue density | -0.48 | -0.33 | 0.16 |
| Root-zone soil pH | 0.13 |  | 0.99 |

